# Comparing Sites of Plasticity in Models of Adaptation to Manifold-Based Perturbations in Brain-Computer Interfaces

**DOI:** 10.1101/2023.03.11.532146

**Authors:** Alexandre Payeur, Amy L. Orsborn, Guillaume Lajoie

## Abstract

During well-trained behaviors, neural population activity in motor cortex lies on a low-dimensional manifold. This raises the question of how such structure constrains subsequent learning. In brain–computer interface experiments in nonhuman primates, perturbations aligned with this subspace induced rapid adaptation, whereas misaligned perturbations induced slower adaptation. Several theoretical accounts have been proposed to explain this differential adaptation, differing in the locus of plasticity. We compare these hypotheses using a minimal linear recurrent network operating at its fixed point and trained by gradient descent. All candidate plasticity sites are able to produce some degree of differential adaptation, whose strength depends on the variance of recurrent weights, with different sensitivities across sites. Hessian analysis reveals how misaligned perturbations reshape the loss landscape by introducing directions of shallow curvature along which gradient descent proceeds slowly. We further propose an experimental test to help distinguish the contributions of different plasticity sites during adaptation. Overall, our results identify the variance of recurrent weights as a key control parameter governing differential adaptation, alongside the site of plasticity.

## 1 Introduction

During skilled behavior, neural population activity often evolves along low-dimensional manifolds embedded in neural space (Gallego, Perich, Miller, & Solla, 2017; Jazayeri & Afraz, 2017; Chung & Abbott, 2021; Langdon, Genkin, & Engel, 2023; Perich, Narain, & Gallego, 2025). These manifolds have been proposed to constrain (Sadtler et al., 2014), and serve as a basis for (Oby et al., 2019), subsequent learning involving the same neural population. Brain–computer interfaces (BCIs) provide experimental access to these constraints by imposing a direct mapping between neural activity and behavior (M. D. Golub, Chase, Batista, & Yu, 2016; Orsborn & Pesaran, 2017). In this setting, the mapping can be constructed so that the low-dimensional latent variables underlying the manifold directly determine the behavioral output, enabling controlled perturbations aligned or misaligned with this structure. In a landmark study, Sadtler et al. (2014) showed that within-manifold (WM) perturbations evoked rapid single-session adaptation, whereas adaptation to outside-manifold (OM) perturbations was limited over the same timescale, according to their performance metrics. Explaining this asymmetry requires identifying how manifold structure interacts with underlying plasticity mechanisms.

Recent theoretical accounts have sought to address this question using recurrent neural network (RNN) models. A key distinction among these models concerns the locus of plasticity: whether adaptation is attributed to recurrent connections (Feulner & Clopath, 2021; Wärnberg & Kumar, 2019), to feedback or input weights (Gurnani, Liu, & Brunton, 2024), or to modifications to the inputs themselves (Menéndez et al., 2024), effectively implementing the reaiming strategy observed in BCI experiments (Jarosiewicz et al., 2008; Chase, Schwartz, & Kass, 2010; Hwang, Bailey, & Andersen, 2013). However, these assumptions have largely been examined in distinct modeling settings, making their relative contributions difficult to compare directly. Here we systematically evaluate these candidate plasticity sites within a unified and minimal framework, and identify the conditions under which each mechanism gives rise to differential adaptation. To isolate the contribution of each mechanism, we study each candidate plasticity site individually, and in the Discussion consider the circumstances under which they could coexist. We further show how perturbations shape the geometry of the loss landscape, influencing learning speed.

To formalize adaptation to manifold-based perturbations, we develop a RNN model that captures the low-dimensional structure of neural activity and its mapping to behavior. To enable analytical insight, we consider linear networks and analyze their fixed-point dynamics. Within this framework, gradient descent serves as a normative proxy (Richards & Kording, 2023) for plasticity targeted at specific circuit components. We find that reaiming produces differential adaptation across parameter regimes, whereas input- and recurrent-weight plasticity do so only when the variance of recurrent weights is sufficiently large, a qualitative distinction that persists in nonlinear networks. In this parameter regime, plasticity can thus be more widely distributed, while still being compatible with differential adaptation. Our framework predicts that evaluating reassociation (M. D. Golub et al., 2018) beyond the latent variables controlling the cursor could help distinguish the contribution of reaiming from other plasticity sites during WM adaptation.

## 2 Methods

Our model captures the key features of the BCI experiment presented in (Sadtler et al., 2014), with targeted simplifications that streamline the analysis. In this experiment, neural activity was recorded from the primary motor cortex of non-human primates using invasive neural implants. This activity was decoded to control a cursor from a center position on a monitor to one of *K* = 8 peripheral targets evenly distributed on a circle, a configuration corresponding to the common center-out paradigm used in BCI research. A central feature of the Sadtler et al. experiment is that the cursor movement is driven by low-dimensional latent factors—defining the manifold—rather than by the full neural activity.

We first introduce the network model, followed by the BCI decoder mapping neural activity to cursor position via the manifold. We then define the WM and OM perturbations and the gradient-based learning rules governing adaptation. Finally, we describe additional analyses used to characterize learning dynamics and the resulting changes in neural activity.

### 2.1 Network model

We model motor cortical activity using a RNN with *N*_tot_ units evolving as

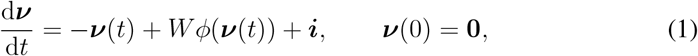

where 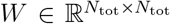 is the recurrent weight matrix and *ϕ* is an activation function. The external drive ***i*** is defined as

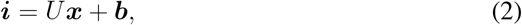

where 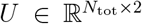 is the input weight matrix and 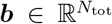 is a bias. The input ***x*** ∈ ℝ^2^ represents static or slowly varying positional information used to reach the current target, and is thus treated as constant within a trial.

To enable learning of target-dependent inputs, we define

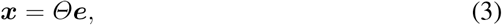

where ***e*** is a one-hot vector representing target identity. This constitutes an idealized representation of task-related inputs upstream of motor cortex, analogous to the abstract command variables considered in (Menéndez et al., 2024). For *K* targets, ***e***^(*k*)^ denotes the *k*th standard basis vector in ℝ^*K*^, and *Θ* ∈ ℝ^2×*K*^ is a learnable matrix parameter. We consider *K* = 8 targets equally spaced on the unit circle,

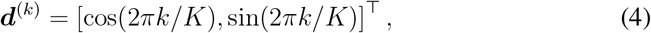

with *k* = 0, …, *K* − 1. At network initialization (denoted by subscript 0), the inputs are identified with the targets,

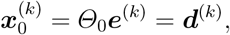

which implies that the columns of *Θ*_0_ are given by the target vectors ***d***^(*k*)^ (see Section 2.5 for more details on network initialization).

Our formulation exposes three key plasticity sites: the inputs (*Θ, reaiming*), the input weights (*U, input-weight learning*), and the recurrent weights (*W, recurrent-weight learning*). The output weights typically present in RNN models are replaced by the BCI decoder described below (Section 2.2); hence, output weights are not among the plasticity sites.

We focus on the linear regime *ϕ*(***ν***) = ***ν***, a common approximation when modeling the motor cortex (Hennequin, Vogels, & Gerstner, 2014; Lara, Cunningham, & Churchland, 2018; Logiaco, Abbott, & Escola, 2021). When *I* − *W* is invertible^1^ and the dynamics are stable, ***ν***(*t*) approaches the fixed point

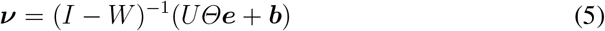

as *t* → ∞. In what follows, we restrict attention to this steady-state response to simplify the analysis.

### 2.2 Manifold and decoder

Following prior theoretical studies, we adopt simplifying assumptions in the BCI de-coder, most notably replacing factor analysis with principal component analysis to define the manifold (Feulner & Clopath, 2021; Gurnani et al., 2024; Menéndez et al., 2024). To avoid introducing a temporary output weight matrix, the manifold is defined at network initialization (Menéndez et al., 2024).

The readout activity ***r*** ∈ ℝ^*N*^ is obtained from the network activity via a *N* × *N*_tot_ measurement matrix *M*,

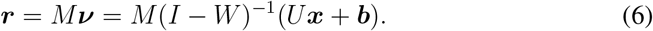

Each row of *M* contains a single entry equal to 1, selecting the corresponding network unit. More general measurement schemes are possible (Menéndez et al., 2024), but we consider only one unit per readout. In the experiments, animals undergo extensive training prior to the application of perturbations (M. D. Golub et al., 2018). To reflect this, readout units are defined as the network units exhibiting the highest variance at initialization, which is intended to partially capture the resulting targeted modulation of readout responses (Ganguly, Dimitrov, Wallis, & Carmena, 2011; Zippi, You, Ganguly, & Carmena, 2022; Rajeswaran, Payeur, Lajoie, & Orsborn, 2025).

The manifold is obtained by diagonalizing the normalized covariance matrix of the readout activity at initialization. Since ***x***_0_ ≡ ***d***, ⟨***x***_0_⟩ = **0**, where 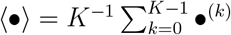. The mean readout activity is thus

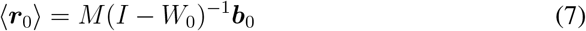

and the covariance matrix is

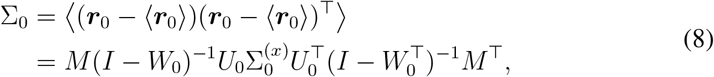

where

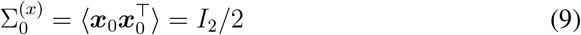

is the input covariance. The normalized covariance is

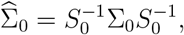

where *S*_0_ is diagonal with 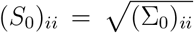. Since 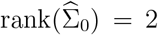, the manifold is two-dimensional and captures all variance. Its basis consists of the two eigenvectors corresponding to the nonzero eigenvalues.

Following experimental procedures, the factors ***z*** ∈ ℝ^2^ used for BCI control are obtained by z-scoring the readout activity, projecting onto the manifold, and z-scoring the result. Defining *E* ∈ ℝ^*N* ×2^ as the reduced eigenvector matrix and Λ ∈ ℝ^2×2^ as the diagonal matrix of nonzero eigenvalues,

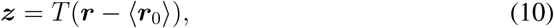

where

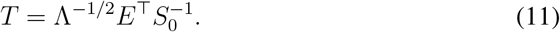

This ensures

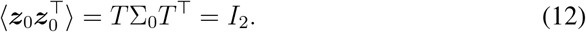

The z-scored readout activity is 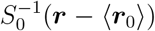, *E*^⊤^ projects it onto the manifold, and Λ^−1/2^ performs the second z-scoring.

The output ***y*** is given by the linear-regression model (“intuitive mapping”)

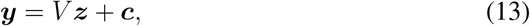

where *V* ∈ ℝ^2×2^ and ***c*** ∈ ℝ^2^ are fitted using ***z*** = ***z***_0_ and ***y*** = ***d***. Since ⟨***z***_0_⟩ = **0** and ⟨***d***⟩ = **0**, we have ***c*** = ⟨***d***⟩ = **0**. The least-square solution is

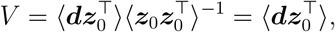

where the last equality uses Eq. 12. From the definition of ***z*** (Eq. 10) together with Eqs. 6 and 7,

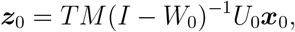

where ***x***_0_ ≡ ***d*** at initialization. Using Eq. 9, we therefore obtain

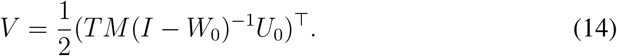

Alternatively, since *V* also satisfies ***d*** = *V* (*TM* (*I* − *W*_0_)^−1^*U*_0_)***d*** at initialization, we can write

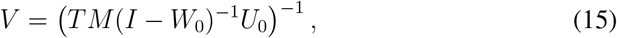

because the inverse exists. Since an exact solution exists, there is no need to train the network after fitting the decoder, because ***y*** ≡ ***d*** already holds. This is a simplification relative to the experiments, in which the animals had to practice with the intuitive decoder during a baseline block before the perturbation was applied (Sadtler et al., 2014). Another consequence is that the least-square solution (Eq. 14) is actually exact (the least square is zero), allowing us to combine both expressions for *V* and write

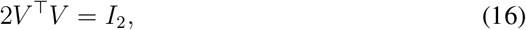

which will be used in Section 3.2.

### 2.3 Perturbations and performance criterion

Within-manifold (WM) perturbations are defined by

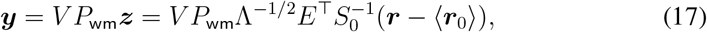

where *P*_wm_ is a 2 × 2 permutation matrix interchanging the components of ***z***. Outside-manifold (OM) perturbations are defined by

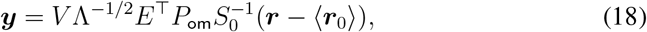

where the *N* × *N* matrix *P*_om_ permutes the components of the z-scored readout activity 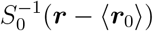. We define

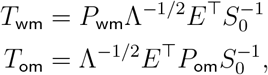

so that during adaptation

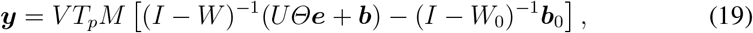

for *p* ∈ {wm, om}.

After a perturbation has been applied, the network is trained to minimize the mean squared error loss

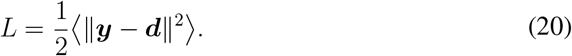

For our two-dimensional manifold, there is a single WM perturbation, corresponding to swapping the rows of *T* . In contrast, there are *N* ! − 1 OM perturbations, which is intractable for the large *N* ∼ 100 used experimentally. Here, we set *N* = 8 readout units, allowing evaluation of all 40,319 nontrivial OM perturbations with reasonable compute time. The ratio of readout units to manifold dimension is 8 : 2 (vs. ∼ 9 : 1 experimentally). To control initial performance impairment (Sadtler et al., 2014), we only analyze OM perturbations whose immediate post-perturbation loss matches that of the WM perturbation within a tolerance (Figure 1A); we refer to these as *valid OM perturbations*.

**Figure 1.**
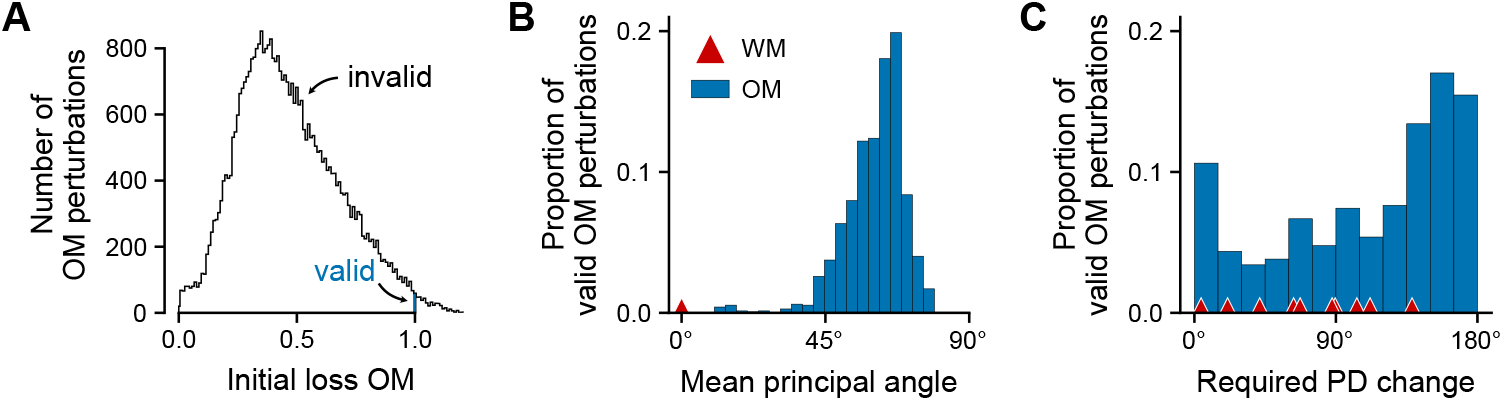
Relative difficulty of perturbations. (A) Initial (pre-adaptation) losses for all *N* ! − 1 OM perturbations for a single random network initialization (black histogram profile). Valid OM perturbations—defined as those with initial losses close to the WM perturbation—are shown in blue. (B) Mean principal angles between intuitive and perturbed mappings (Eq. 31) for valid OM perturbations, across *n* = 10 network initializations. Red triangles indicate mean principal angles for WM perturbations, which are overlapping here. (C) Mean required change in preferred direction (PD); same format 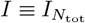.

### 2.4 Learning

We derive the gradient-descent learning dynamics using three standard identities: 1) for a scalar function of a matrix variable *f* (*X*), th_}_e differential is d*f* = tr{∇_*X*_*f* ^⊤^d*X*} ; 2) the trace tr is cyclic; and 3) ***f*** ^⊤^***g*** = tr {***gf*** ^⊤^} for any vectors ***f*** and ***g***. Defining the error as ***ϵ*** = ***y*** − ***d***, the differential of the loss is

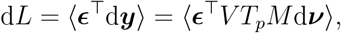

where d***ν*** is the differential of Eq. 5. Using

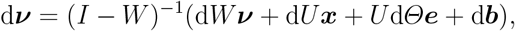

we obtain, after straightforward manipulations,

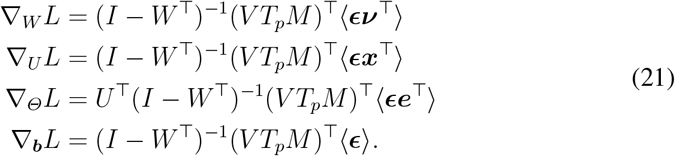

It is convenient to rewrite these gradients in matrix form. For any two vectors 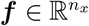 and ***h*** ∈ 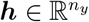 taking *K* values,

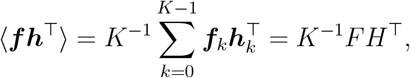

where *F* = [***f***_0_, …, ***f***_*K*−1_] is a *n*_*x*_ × *K* matrix and similarly for *H*. We define the output matrix *Y* = [***y***_0_, …, ***y***_*K*−1_], the target matrix *D* = [***d***_0_, …, ***d***_*K*−1_], the error matrix ℰ = *Y* − *D* and the *N*_tot_ × *K* neural activity matrix *N* = [***ν***_0_, …, ***ν***_*K*−1_]. We also define the broadcasted bias matrix 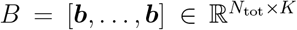. With these definitions,

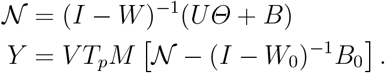

The gradients with respect to the matrix parameters can then be written as

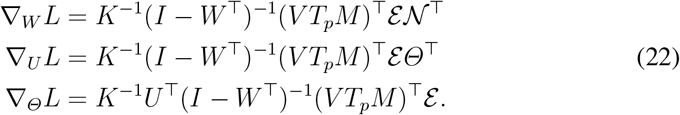

Parameters are updated according to

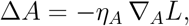

where *A* ∈ {*W, U, Θ*, ***b*}** and *η*_*A*_ is the learning rate. To ensure comparable performances across network initializations and learning mechanisms, *η*_*A*_ is selected via a log-scale bisection search such that the final loss after WM adaptation (evaluated at epoch 150, i.e., 1200 trials, comparable to experiments) falls within a prescribed range (e.g., [10^−4^, 5 × 10^−4^]), corresponding to near-complete adaptation.

### 2.5 Initialization

Recurrent weights were initialized as 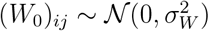. By Girko’s circular law, the spectral radius of *W* —the largest eigenvalue magnitude—scales as 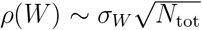 for large *N*_tot_. We therefore choose 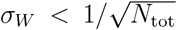 to promote stability. At finite *N*_tot_, however, eigenvalue outliers may occur (Rajan & Abbott, 2006). Moreover, when *W* is learned, this condition does not in general guarantee stability during adaptation (Section 3.4). To mitigate instabilities, we set the largest *σ*_*W*_ to 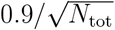 and monitor the spectral radius throughout training.

Input weights were initialized as 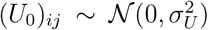. The choice of 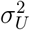 has only a marginal impact on adaptation for two reasons. First, the network output ***y*** at initialization is invariant under rescaling of *U*_0_, i.e., 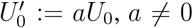, because the decoder compensates for this change. Second, *U*_0_ enters linearly into the learning rules (Eq. 22), so any rescaling can be absorbed into the learning rate. When *U* is the learnable parameter, we verified that our results are largely insensitive to *σ*_*U*_ (Figure 5A).

**Figure 2.**
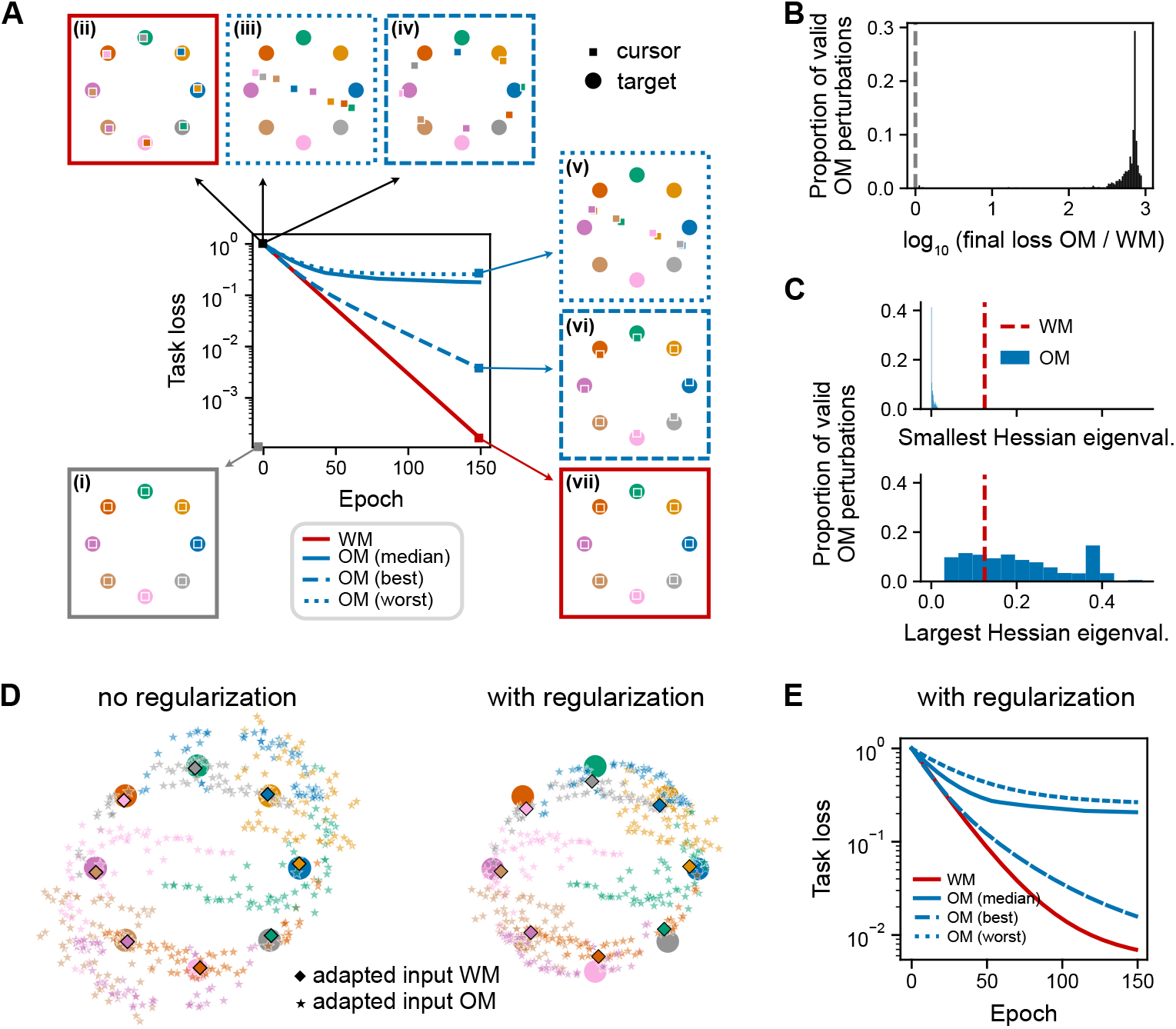
Reaiming. (A) Task loss during adaptation for a selected random network initialization. One epoch corresponds to reaching to all targets. (i-vii) Model outputs (squares) and targets (circles) at successive stages: (i) pre-perturbation; (ii-iv) post-perturbation, pre-adaptation; (v-vii) post-adaptation. OM training runs with the largest (worst, dotted) and smallest (best, dashed) final losses are highlighted. (B) Distribution across random initializations (*n* = 10) of the log_10_ ratio of final task losses (OM/WM) for valid OM perturbations. (C) Distribution of smallest (top) and largest (top) eigen-values of the Hessian matrix, across seeds. (D) Learned inputs (columns of *Θ*^∗^) for the same network initialization as in panel A, without (*λ* = 0; left) and with regularization (*λ* = 2 × 10^−4^; right). (E) Task loss with regularization (same initialization as in panel A).

**Figure 3.**
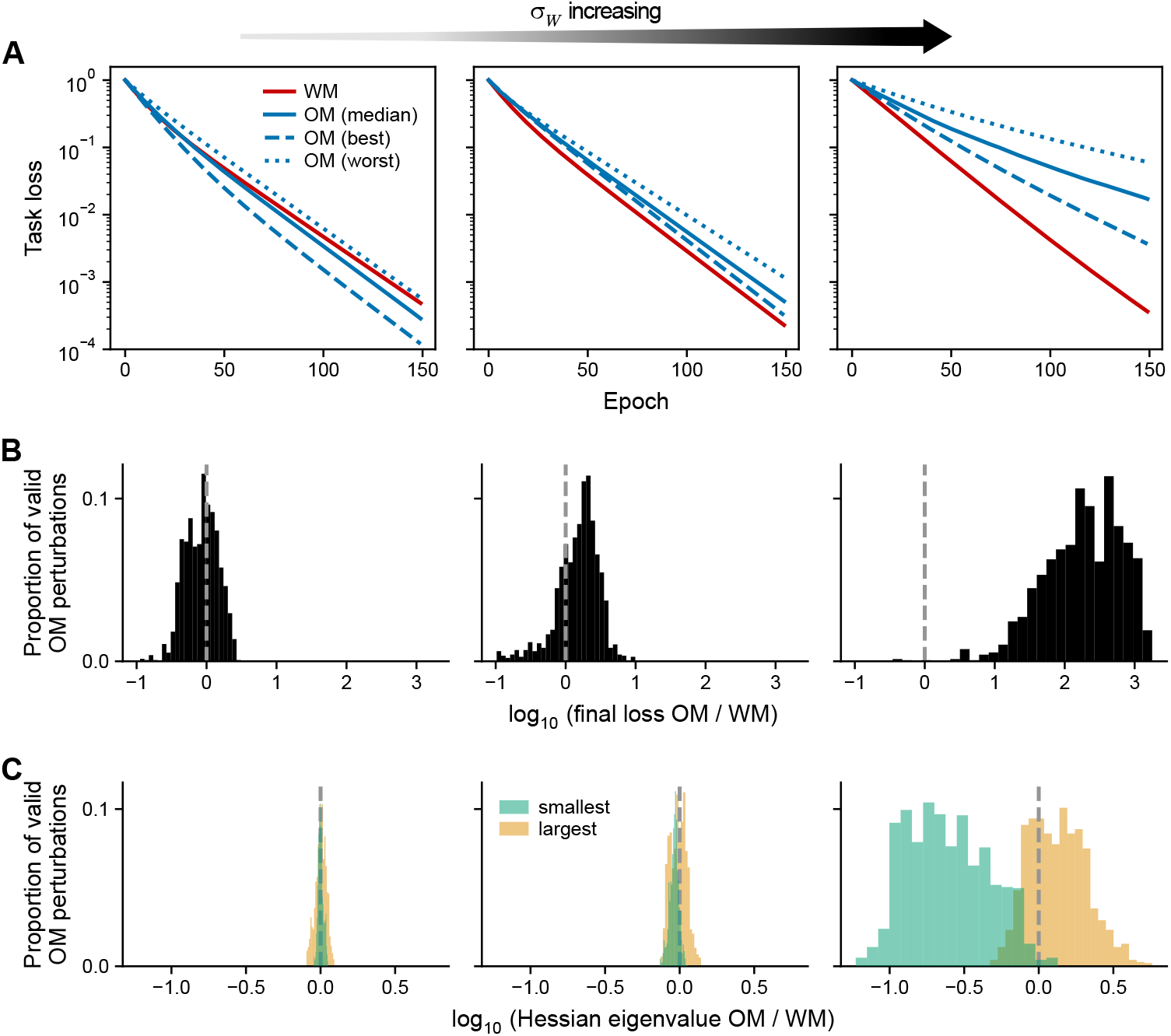
Input-weight learning. In all panels, the recurrent-weight standard deviation *σ*_*W*_ increases from left to right: left: 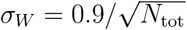; center: 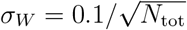; right: 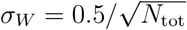. (A) Task loss during adaptation for a selected network initialization. (B) Distribution across random initializations (*n* = 10) of the log ratio of final task losses (OM/WM) for valid OM perturbations. (C) Distribution of the log ratio of the smallest and largest eigenvalues of the Hessian (OM/WM).

**Figure 4.**
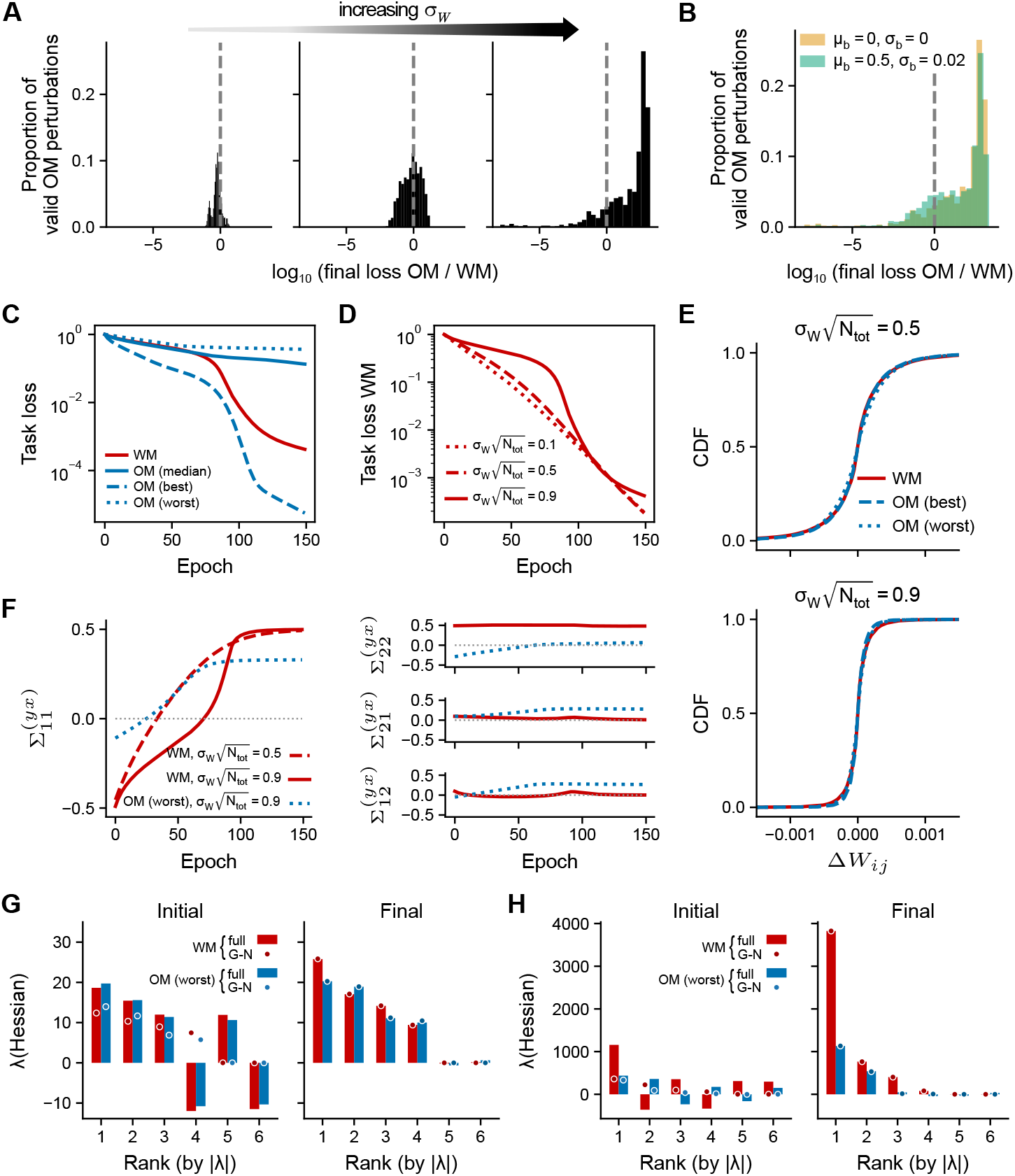
Recurrent-weight learning. (A) Distribution across random initializations (*n* = 10) of the log ratio of final task losses (OM/WM) for valid OM perturbations for 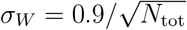 (left), 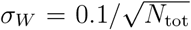 (center), 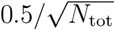. (B) Same as panel A, right, for ***b***_0_ = **0** and 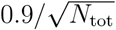. Panels C-H refer to the same representative network initialization. (C) Task loss (same format as in Figure 2A). (D) Task loss for the WM perturbation for different *σ*_*W*_ . (E) Cumulative density function of the change in recurrent weight, Δ*W*_*ij*_ = *W*_*ij*_ − *W*_0,*ij*_ across all *ij* pairs, accumulated during adaptation. (F) Input-output covariance entries for the model. (G) Top-6 Hessian eigenvalues ranked by their absolute values for 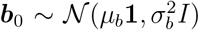, before (left) and after (right) adaptation. G-N: Gauss-Newton. (H) Same as G, for 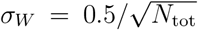.

**Figure 5.**
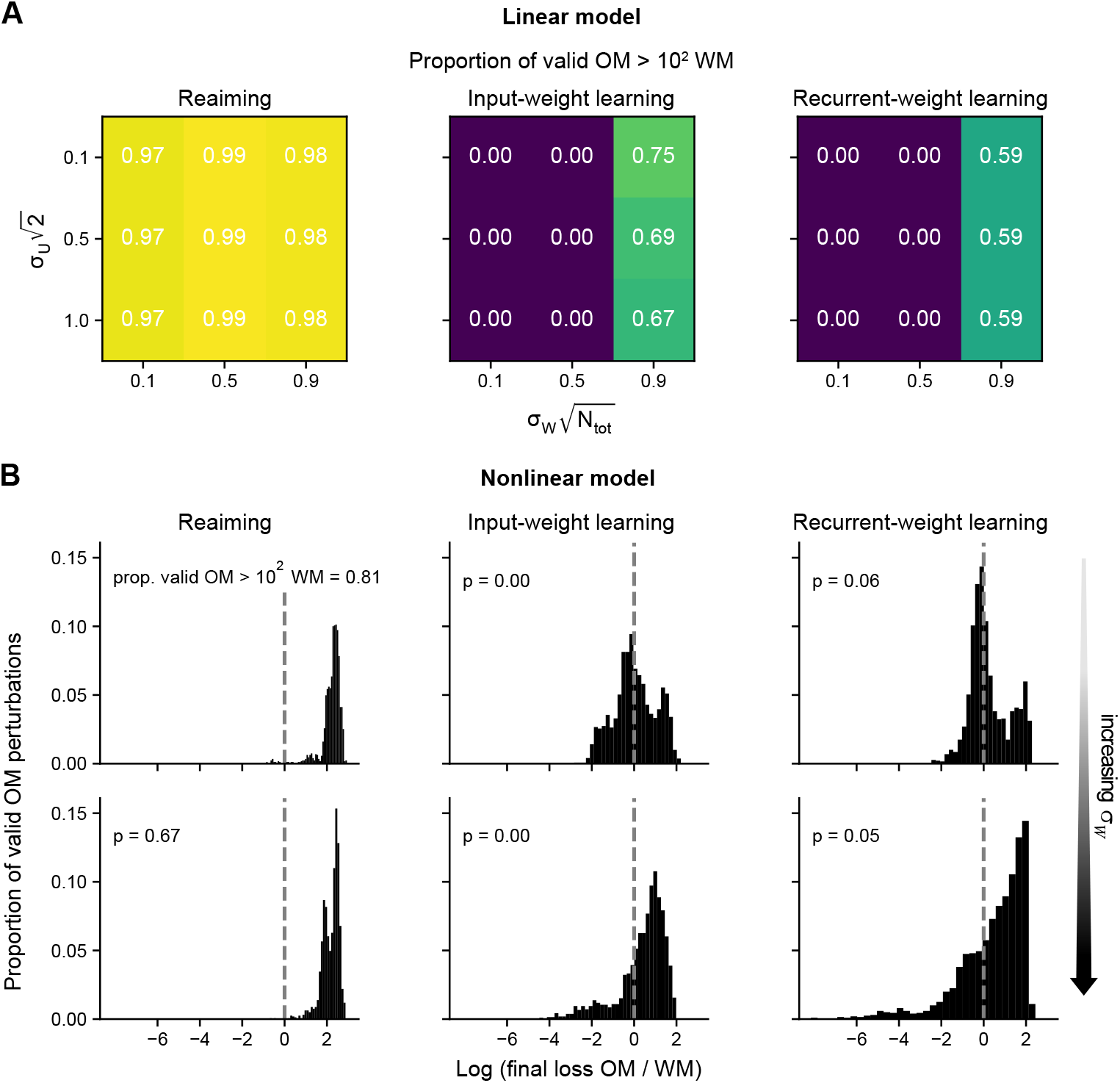
Extension to nonlinear RNNs. (A) Proportion of valid OM perturbations with final loss greater than 10^2^ times the final WM loss as a function of *σ*_*U*_ and *σ*_*W*_ . (B) Distribution across random initializations (*n* = 20) of the log_10_ ratio of final task losses (OM/WM) for valid OM perturbations. Parameters: ***b***_0_ = **0**, 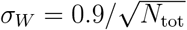 (top) and 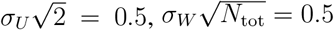 (bottom).

The initialization of the bias ***b*** also plays a marginal role. Because BCI control uses z-scored activity, the contribution of the bias to the output is eliminated whenever *W* was not learned (Eq. 19). The case in which *W* is learned is discussed in Section 3.4.

Unless stated otherwise, we used 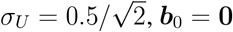 and *N*_tot_ = 500.

### 2.6 Hessian

The Hessian characterizes the local curvature of the loss landscape and is therefore in-formative for analyzing learning dynamics. Using the error matrix ℰ = *Y* − *D* (Section 2.4), the loss can be written as

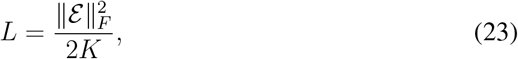

where ∥ • ∥_*F*_ denotes the Frobenius norm. The first-order differential is

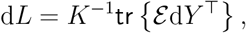

and the second-order differential is

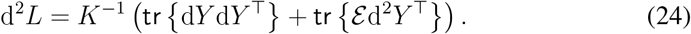

The first term is the Gauss-Newton contribution.

#### Plastic *Θ*

When *Θ* is the only learnable parameter,

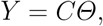

where *C* is a constant 2 × 2 matrix defined by

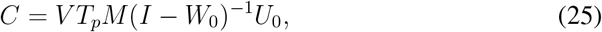

with *p* ∈ {wm, om} . For brevity, we omit the subscript *p* on *C* and on all quantities that depend on *T*_*p*_. Then d*Y* = *C* d*Θ* and d^2^*Y* = 0, so

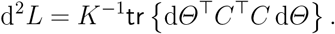

Vectorizing with the identities

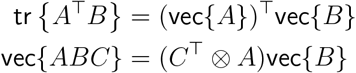

for arbitrary matrices *A, B, C* (Magnus & Neudecker, 2019), *we obtain*

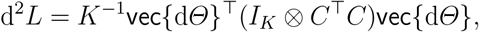

where ⊗ denotes the Kronecker product. Hence, the 2*K* × 2*K* Hessian is

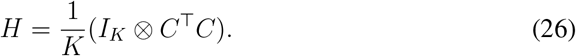

#### Plastic *U*

When *U* is the only learnable parameter,

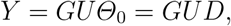

where *G* is a constant 2 × *N*_tot_ matrix defined by

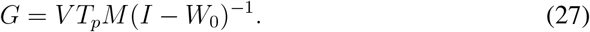

Using d*Y* = *G* d*UD*, d^2^*Y* = 0, and *DD*^⊤^ = (*K*/2)*I*_2_, we obtain

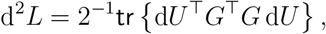

and the 2*N*_tot_ × 2*N*_tot_ Hessian matrix is

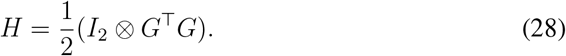

#### Plastic *W*

When *W* is the only learnable parameter (assuming ***b***_0_ = **0**),

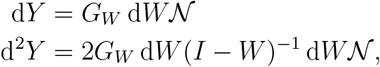

where

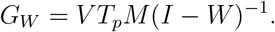

The first term in Eq. 24 becomes

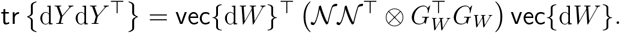

The second term involves d*W* twice (rather than d*W* and d*W* ^⊤^). To handle this, we use the commutation matrix *K*, defined by *K*vec{d*W* ^⊤^} = vec{d*W*} (Magnus & Neudecker, 2019). Since *K* is orthogonal, *K*vec {d*W* ^⊤^} = (vec{d*W* ^⊤^})^⊤^, which yields

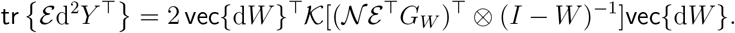

This contribution must be symmetrized. The resulting 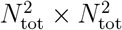 Hessian is

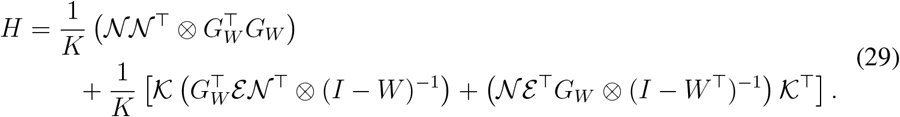

Unlike the Hessians for *Θ* and *U*, the Hessian for *W* evolves during learning.

### 2.7 Procrustes distance

We denote generic pre- and post-adaptation activity matrices by *X*_pre_ and *X*_post_, both of size *n* × *K* matrices (with *n* = 2, *N*, or *N*_tot_ depending on the activity type). Their similarity was quantified using the Procrustes distance

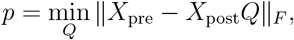

subject to *Q*^⊤^*Q* = *I*_*K*_. If *X*_post_ differs from *X*_pre_ only by an orthogonal transformation— including rotations and permutations of target labels—then *p* = 0. Equivalently (G. H. Golub & Van Loan, 2013),

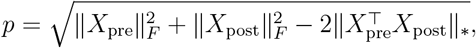

where ∥ • ∥_∗_ denotes the nuclear norm. If *X*_post_ = *αX*_pre_, then *d* = |1 − *α*|∥*X*_pre_∥_*F*_, motivating the normalized distance

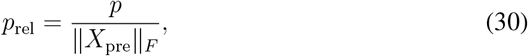

which measures fractional changes relative to the pre-adaptation condition and accounts for differences across activity types and random network initializations.

## 3 Results

We first assess the relative difficulty of WM and OM perturbations in the model and compare them with experimental results. Next, we isolate each learning parameter’s contribution to adaptation by allowing plasticity in only one parameter at a time^2^, and then test whether these observations extend to nonlinear activation functions. Finally, we analyze the neural strategies underlying WM adaptation across the different plasticity mechanisms.

### 3.1 Relative difficulty of perturbations

In (Sadtler et al., 2014), several controls ensured that OM perturbations were not artificially more difficult to learn due to trivial differences in scaling or effective search-space dimensionality. We adopted analogous precautions. Neural activity and latent factors were z-scored, and we exhaustively searched for OM perturbations whose pre-adaptation losses lay within a specified tolerance of that of the WM perturbation, which we call valid OM perturbations (Figure 1A). Consequently, the initial performance impairments were comparable across perturbation types.

Sadtler et al. (2014) also performed post hoc controls to assess the expected relative difficulty of adaptation. One such measure is the mean principal angle between the intuitive and perturbed mappings, which quantifies the deviation between their row spaces.

These mappings connect the z-scored readout activity, *S*^−1^(***r*** − ⟨***r***_0_⟩), to the output ***y*** in Eqs. 13, 17, and 18:

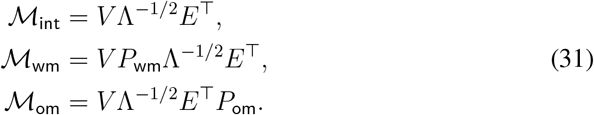

Each is a 2 × *N* matrix whose rows span a two-dimensional subspace of ℝ^*N*^, yielding two principal angles. We report their mean for WM and OM perturbations, denoted ∠( ℳ_int_, ℳ_wm_) and ∠(ℳ_int_, ℳ_om_), respectively (Figure 1B). For WM perturbations, the mean principal angle is identically zero across all network initializations (Figure 1B, overlapping red triangles). This is because *V* is invertible and *P*_wm_ implements a row interchange; both operations preserve the row space. In contrast, valid OM perturbations yield a distribution of mean principal angles (60^°^ 10^°^, mean s.d.) across network initializations. This discrepancy with experiments arises from the two-dimensional manifold which captures 100% of the variance. Importantly, as shown in Sections 3.3-3.4, large principal angles do not necessarily imply slower adaptation: for some parameter regimes, adaptation rates are comparable across perturbation types.

Another control considered by Sadtler et al. is the expected change in preferred direction (PD) induced by the perturbations. Each of the *N* columns of the intuitive mapping in Eq. 31—one per readout unit—represents a “pushing” direction in the two-dimensional workspace. Perturbations alter these directions, requiring coordinated changes in read-out activity to generate the correct output. Accordingly, the change in push direction induced by a perturbation serves as a measure of the required PD change (Figure 1C). In our model, both WM and OM perturbations evoked a wide range of required PD changes, suggesting comparable demands on the readout units.

### 3.2 Reaiming (plastic *Θ*)

We first consider the case in which the input ***x***, parametrized by *Θ*, is the sole learnable quantity. This corresponds to reaiming (Jarosiewicz et al., 2008), later formalized as a general theory of BCI learning (Menéndez et al., 2024).

We specialize standard results on gradient-descent dynamics for linear feedforward neural networks to our setting. Using the definition of *C* (Eq. 25), the error matrix ℰ = *CΘ* − *D* and Eq. 22, the gradient-descent dynamics are

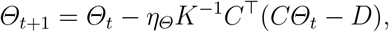

where the subscript *t* denotes the update step. These linear dynamics converge to the unique fixed point *Θ*^∗^ = (*C*^⊤^*C*)^−1^*C*^⊤^*D*, provided that *C*^⊤^*C* is full rank and

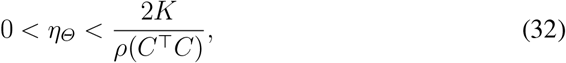

where *ρ*(*C*^⊤^*C*) denotes the spectral radius. When *C* is invertible—this holds across all tested network initializations and valid OM perturbations—the fixed point simplifies to

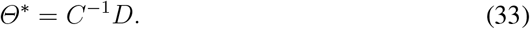

Since *C* is invertible, at convergence ℰ^∗^ = 0 and the loss vanishes regardless of the perturbation; what differs are the relative convergence rates for WM and OM. The error evolves according to

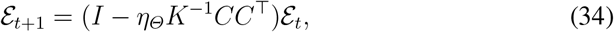

with 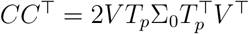 (Eq. 8). For WM, because *P*_wm_ is orthogonal and *T* Σ_0_*T* ^⊤^ = *I*_2_, we have *CC*^⊤^ = 2*V V* ^⊤^ = *I*_2_ (by Eq. 16). From Eqs. 23 and 34, the loss then decays exponentially:

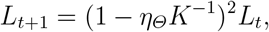

consistent with simulations (Figure 2A). The fixed parameters *W*_0_, *U*_0_, ***b***_0_ therefore play no role in WM adaptation. In contrast, for OM, *CC*^⊤^ ≠ *I*_2_, so the loss dynamics depend on the specific perturbation (Figure 2A) and network initialization. Although zero loss remains achievable, the limited training duration (150 epochs = 1200 trials, matching the experimental setting) and chosen learning rate yield substantially worse final performance for OM than for WM (Figure 2B). Varying the variances of *U* and *W* initialization changes *C* but has only a modest quantitative effect on final performance (Section 3.5, Figure 5A, left).

To understand the difference in adaptation rates, we compute the Hessian of the task loss (Eq. 26). The Hessian is block-diagonal with identical blocks *K*^−1^*C*^⊤^*C*, so its spectrum consists of *K* copies of the two eigenvalues of *K*^−1^*C*^⊤^*C*. For WM, *C*^⊤^*C* = *I*, so the Hessian is isotropic with all eigenvalues equal to *K*^−1^ (Figure 2C). In contrast, OM perturbations introduce a small eigenvalue—corresponding to *K* relatively flat directions in the loss landscape—along which gradient descent proceeds more slowly (Figure 2C).

During a single experimental session, OM adaptation appears impossible, whereas WM adaptation either seemingly plateaus at a suboptimal solution (figure 2a in (Sadtler et al., 2014)) or continues to improve slowly (figure 1c in (M. D. Golub et al., 2018)). In contrast, reaiming in the model can achieve complete adaptation to WM perturbations. The corresponding solution, *Θ*^∗^ = *C*^⊤^*D*, preserves input norms because *C* is orthogonal, whereas OM perturbations can induce deviations from unit norm (Figure 2D, left). To illustrate how a competing objective could limit adaptation, we add an activity regularization term

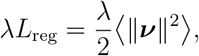

where *λ >* 0 controls the trade-off. At convergence (with ***b***_0_ = **0**), the regularized solution is *Θ*^∗^ = (*C*^⊤^*C* + *λF* ^⊤^*F*)^−1^*C*^⊤^*D*, where *F* = (*I − W*_0_)^−1^*U*_0_, leading to a shrinkage of input norms (Figure 2D, right). With regularization, WM adaptation becomes incomplete, converging to a higher final loss, while OM adaptation is comparatively less affected (Figure 2E). This shows that a competing objective of this form can reproduce incomplete WM adaptation, although activity regularization may not be the exact underlying cause (see Discussion).

### 3.3 Input-weight learning (plastic *U*)

We next consider *input-weight learning*, in which *U* is the sole learnable parameter. The network output is *Y* = *GUD*, with *G* defined in Eq. 27. The gradient ∇_*U*_ *L* = *K*^−1^*G*^⊤^ℰ *D*^⊤^ (Eq. 22) induces linear dynamics

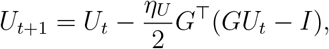

where we used *DD*^⊤^ = (*K*/2)*I*_2_. At convergence, *U* ^∗^ satisfies *G*^⊤^*GU* ^∗^ = *G*^⊤^, an underdetermined system since rank(*G*^⊤^*G*) = 2. Its solution decomposes as

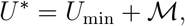

where *U*_min_ = *G*^⊤^(*GG*^⊤^)^−1^ is the minimum-norm solution and ℳ ∈ ker *G*, i.e., *G*ℳ = 0. Writing 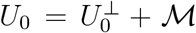 at initialization, with 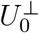 orthogonal to ker{*G*}, gradient descent drives 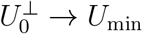^3^ while leaving ℳ unchanged (Appendix B). Thus, only the component of *U* in the range of *G*^⊤^ is shaped by learning. Because *G* differs between WM and OM perturbations (Eq. 27), the accessible component range {*G*^⊤^} changes, and we expect this restriction to differentially affect adaptation.

Interestingly, the emergence of distinct learning curves for WM vs. OM perturbations depends on the variance of the recurrent weights (Figure 3A-B). A noticeable difference appears when *σ*_*W*_ approaches instability; if *σ*_*W*_ is too small, WM and OM perturbations are roughly equally difficult. To understand this, we analyze the Hessian spectrum for input-weight learning (Eq. 28). As in reaiming, the Hessian is fixed during learning and block-diagonal, with each block equal to *G*^⊤^*G*/2 and possessing only two nonzero real eigenvalues (since *G* has rank 2). Consequently, the Hessian spectrum contains four nonzero eigenvalues—two copies of the two nonzero eigenvalues *G*^⊤^*G*/2—and 2(*N*_tot_ − 2) zero eigenvalues. This structure is consistent with the analysis of the solutions at convergence, *U* ^∗^. The loss landscape therefore contains many flat directions, irrespective of *σ*_*W*_ . However, as *σ*_*W*_ increases, the smallest nonzero eigenvalue of the OM Hessian becomes, on average, roughly half an order of magnitude smaller than that of WM (Figure 3C), contributing to the observed differential adaptation.

### 3.4 Recurrent-weight learning (plastic *W*)

We next consider *recurrent-weight learning*, where *W* is the sole learnable parameter. Because adaptation may drive the spectral norm of *W* beyond the instability threshold of 1, we initialize it well below this threshold and monitor it throughout. OM perturbations that cause the spectral norm to exceed 1 are excluded from the set of valid perturbations.

As in input-weight learning, differential adaptation emerges when *σ*_*W*_ is sufficiently large (Figure 4A). In contrast to reaiming and input-weight learning, the constant bias is uncompensated in the error

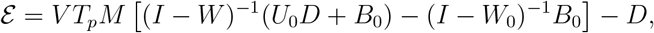

and also appears in the network activity *N* in the loss gradient (Eq. 22). Thus, ***b***_0_ can influence adaptation under recurrent-weight learning. At large *σ*_*W*_, introducing a small heterogeneous bias reduces the difference between WM and OM adaptation (Figure 4B). Larger biases, combined with our learning-rate search procedure, produced adaptation curves that are widely inconsistent across random initializations and often markedly noisy; we therefore do not analyze them further.

When ***b***_0_ = **0**, the loss gradient with respect to *W* is invariant under a rescaling of *U*_0_ (Appendix C). The input variance *σ*_*U*_ can thus be chosen arbitrarily. At high *σ*_*W*_, WM adaptation exhibits a slow initial phase, during which the WM loss remains close to the median OM loss (Figure 4C). This is followed by a sudden decrease in WM loss, which separates from the OM curve. This slow initial phase is absent at smaller *σ*_*W*_ (Figure 4D). Increasing *σ*_*W*_ is associated with smaller changes in recurrent weights during adaptation (Figure 4E). However, as noted in (Feulner & Clopath, 2021), the distribution of weight changes alone does not account for the difference in final performance between WM and OM perturbations (Figure 4E, bottom).

We relate the sudden acceleration of WM adaptation to the alignment between the input-output covariance of the model, denoted Σ^(*yx*)^, with the task covariance, Σ^(*dx*)^ = *K*^−1^*DD*^⊤^ = *I*_2_/2. One diagonal component of Σ^(*yx*)^ changes sign and converges toward the corresponding component of Σ^(*dx*)^ (Figure 4F, left). Meanwhile, the worst OM perturbation saturates below the target value of 1/2, contributing to poor final performance (Figure 4F, left). For WM perturbations, the remaining entries of Σ^(*yx*)^ stay close to their targets throughout adaptation, whereas for the most difficult OM perturbation all entries end far from their respective targets (Figure 4F, right).

To further understand differential adaptation at high *σ*_*W*_, we analyze the Hessian (Eq. 29). In contrast to reaiming and input-weight learning, the Hessian varies throughout adaptation. The Gauss-Newton contribution, 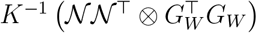, has rank 4, as in input-weight learning, but is not block-diagonal because it depends on the full network covariance (for ***b***_0_ = **0**). Its four nonzero eigenvalues are given by pairwise products of the two nonzero eigenvalues of *K*^−1^ *NN*^⊤^ and 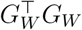. In addition, the Hessian includes error-dependent contributions with rank at most 2*N*_tot_, so the Gauss–Newton spectrum alone does not fully characterize the curvature. We compute the six leading eigenvalues of the full Hessian before and after learning, which appear to capture the main changes in curvature (Figure 4G–H).

Irrespective of *σ*_*W*_, training reduces the effective rank of the Hessian, yielding a largely convex local landscape dominated by the Gauss-Newton component (Figure 4G-H, right). At low *σ*_*W*_, the eigenvalues remain similar for the WM perturbation and the most difficult OM perturbation throughout training (Figure 4G). In contrast, at high *σ*_*W*_, marked differences emerge at the end of training (Figure 4H, right). For the network initialization shown in Figure 4C-H, the nonzero Gauss-Newton eigenvalues are 3843, 769, 427, 85 for WM and 1128, 52, 14, 0.65 for the most difficult OM perturbation. The OM Hessian is therefore highly ill-conditioned (1128/0.65 ≈ 1735), with a slow direction associated with the smallest eigenvalue. As in reaiming and input-weight learning, this suggests that difficult OM perturbations participate in creating slow directions in the loss landscape, limiting adaptation. However, in the recurrent-weight case, the Gauss-Newton eigenvalues also depend on the total covariance, whose eigenvalues all exceed 1, thereby increasing curvature along all directions and partially mitigating this effect.

### 3.5 Extension to nonlinear RNNs

The static linear network allows a systematic characterization of adaptation to manifold-based perturbations. Reaiming is robust to parameter changes: differential adaptation is observed across a wide range of network initializations. In contrast, input-weight and recurrent-weight learning show little differential adaptation unless *σ*_*W*_ is sufficiently large (Figure 5A).

We test whether these observations extend to nonlinear dynamics by replacing the linear activation with a rectified linear unit (ReLU), a standard choice in closely related models (Menéndez et al., 2024; Gurnani et al., 2024). Equation 1 is integrated using the Euler method with initial condition ***ν***_0_ = **0**. When stable, the dynamics converge to an input-dependent fixed point at which the objective (Eq. 20) is evaluated and gradient descent is applied; we use *T* = 40 time steps. The nonlinear setting introduces additional constraints. Learning is less stable, requiring a higher target WM loss to ensure a successful learning-rate search. Moreover, valid OM perturbations are not obtained for all initializations; we therefore use 20 seeds, yielding 11 ± 2 (mean ± sd) successful initializations across learning conditions. Finally, because the ReLU nonlinearity can effectively silence units, we set 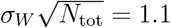 in the high-variance condition.

As in the linear case, reaiming remains largely insensitive to *σ*_*W*_, while the distributions of log ratios of OM and WM final losses for input-weight and recurrent-weight learning become increasingly left-skewed as *σ*_*W*_ increases. However, the proportion of valid OM perturbations whose final loss exceeds 10^2^ times the WM final loss—using the same threshold as in Figure 5A—drops to near zero for input-weight learning and recurrent-weight learning (Figure 5B). Although this threshold is somewhat arbitrary, these results indicate that differential adaptation at these plasticity sites is reduced in the ReLU case compared to the linear case. Overall, the main insights from the linear model are preserved in this nonlinear case, albeit with attenuated quantitative separation.

### 3.6 Reassociation

Reassociation is a learning strategy hypothesized to occur following a WM perturbation, whereby post-training population activity patterns resemble those before training (M. D. Golub et al., 2018). Importantly, *population activity patterns* refer to the latent factors ***z*** that drive BCI control, rather than the readout activity itself (M. D. Golub et al., 2018).

To obtain a comprehensive picture of repertoire changes during WM adaptation in the linear model (Eq. 5), we used the relative Procrustes distance (*p*_rel_, Eq. 30) to quantify the similarity between the pre- and post-training latent factors ***z*** (Eq. 10), readout activity ***r*** and network activity ***ν***. Notably, reassociation corresponds to small values of *p*_rel_. We use a high *σ*_*W*_ so that input-weight and recurrent-weight learning operate in a regime exhibiting differential adaptation (Figure 5A).

As expected, reaiming is fully compatible with reassociation (Menéndez et al., 2024) (Figure 6). Surprisingly, for the other learning mechanisms, the relative distance for the latent factors remains close to zero, indicating that they are also consistent with reassociation (Figure 6A). In contrast, for input-weight learning and recurrent-weight learning, *p*_rel_ deviates from zero for the readout and network activity (Figure 6B-C). This deviation is significantly larger for recurrent-weight learning than for input-weight learning (***r***: *p* = 2 × 10^−3^, ***ν***: *p* = 1 × 10^−3^; Wilcoxon signed-rank test, *n* = 10). Thus, while all learning mechanisms are compatible with reassociation at the level of the latent factors ***z***, clear differences appear at the level of readout and network activity.

**Figure 6.**
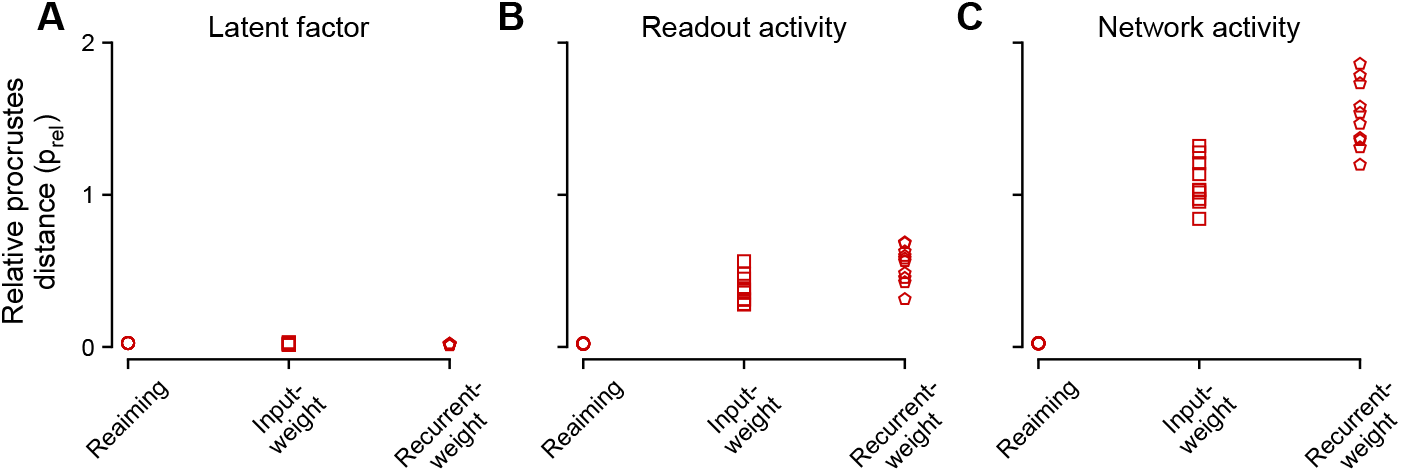
Reassociation. Relative Procrustes distance (*p*_rel_) between pre- and post-training latent factors (A), readout activity (B), and network activity (C), across plasticity sites. Distances are near zero for latent factors across all plasticity sites, consistent with reassociation. Parameter: 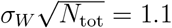.

## Discussion

Our study compares different hypotheses about how pre-existing neural structures may constrain subsequent learning, using the site of plasticity as the key distinguishing factor. We find that differential adaptation can emerge across all considered plasticity sites, although the conditions under which it occurs depend on the site. Reaiming robustly produces differential adaptation across parameter regimes. In contrast, input-weight and recurrent-weight learning require a large recurrent-weight variance *σ*_*W*_, placing the linear network near instability. These findings largely persist qualitatively in networks with nonlinear activation functions. Mechanistically, difficult OM perturbations induce shallow directions—associated with small Hessian eigenvalues—in the loss landscape, along which gradient descent proceeds slowly. Differential adaptation under reaiming is less sensitive to *σ*_*W*_ because the Hessian for WM adaptation is isotropic with large eigenvalues irrespective of the network’s operating point.

After a perturbation, the network must generate activity that is both output-potent (Kaufman, Churchland, Ryu, & Shenoy, 2014) and compatible with the task objective. Previous theories have assumed that the local recurrent connectivity is not plastic on the time scale of the experiment (Sadtler et al., 2014; Zhou, Tien, Ravikumar, & Chase, 2019), and approached this problem by viewing the recurrent network together with the perturbed decoder as a controlled dynamical system (Menéndez et al., 2024; Gurnani et al., 2024). Adaptation becomes an open-loop (Menéndez et al., 2024) and/or feedback (Gurnani et al., 2024) control problem. In these frameworks, learning is constrained by the assumed low dimensionality of the control inputs. Such constraints limit the controllability of network states, particularly when the input magnitude is penalized (Menéndez et al., 2024) or when perturbations require substantial reorganization of pre-existing dynamical flows (Gurnani et al., 2024; Oby et al., 2025). Our results complement these perspectives by showing how perturbations directly reshape the optimization landscape associated with reaiming and input-weight learning. Activity regularization plays a role similar to input-norm constraints by competing with the task objective and limiting WM performance. Although reaiming has been modeled as an open-loop mechanism—in (Menéndez et al., 2024) and here—extensions to feedback control are possible (Menéndez et al., 2024), suggesting potential links with Gurnani et al.’s frame-work.

In these control-theoretic accounts, low-dimensional inputs play a central role in con-straining both adaptation and the intrinsic dimensionality of neural activity. However, the origin of this low-dimensional structure is typically left unspecified and merely reflects the task structure. One may instead embed the latent structure directly into the network architecture (Wärnberg & Kumar, 2019), but then learning mechanisms compatible with this structure may be lacking (Feulner & Clopath, 2021). Another possibility is that adaptation relies less on low-dimensional external inputs and more on recurrent mechanisms, since small correlated changes to local connectivity may suffice for adaptation (Feulner et al., 2022), especially at high *σ*_*W*_ . However, our analysis suggests that recurrent-weight learning early in adaptation is associated with a complex, non-convex loss landscape and slow adaptation rates at high *σ*_*W*_ . Gurnani et al. also articulate cogent arguments against it. A solution explored in (Feulner & Clopath, 2021) was to consider imperfect credit assignment during recurrent-weight learning, but it is unclear how adaptation rates are impacted, especially early during adaptation. Beyond BCI research, fast adaptation to visuomotor perturbations in arm-reaching tasks is associated with minimal changes in motor-cortical functional connectivity, potentially arguing against recurrent-weight learning as a mechanism for rapid learning (Perich, Gallego, & Miller, 2018).

All theoretical accounts of Sadtler et al.’s experiments—including the present work— have relied on modeling choices that can differ from the experimental setting. A few of these choices might deserve reexamination in future work, specifically concerning manifold-based perturbations, but also center-out BCI experiments more generally. A key discrepancy is that, in actual center-out BCI tasks, animals must typically reach the target before a set time interval expires to get rewarded. This challenges the use of a mean-square loss at a precise end time as the primary objective and, more generally, the use of supervised-learning methods instead of reinforcement-learning ones in motor control (Codol, Krishna, Lajoie, & Perich, 2024). Related to this point, viewing adaptation simply as an optimization process might disregard important confounding factors (Hennig et al., 2021). One such factor that has not been widely discussed is that, in perturbative experiments, the animals have had extensive training with the task prior to the perturbation (M. D. Golub et al., 2018) (which we partially accounted for by using the most modulated readout units for control). Highly-trained subjects have exhaustively sampled neural and task variability structures. Structure learning (or meta-learning) could thus speed up adaptation when perturbations share sufficient statistical structure with prior exposure (Braun, Aertsen, Wolpert, & Mehring, 2009; Braun, Mehring, & Wolpert, 2010; Chang, Perich, Miller, Gallego, & Clopath, 2024), without requiring substantial plasticity (Weinstein & Botvinick, 2017). However, it is unclear how such meta-learning processes would interact with the perturbation-induced changes to the optimization landscape identified here.

Within our framework, reaiming is the most robust mechanism for producing differential adaptation, as it does so across parameter regimes and even in the presence of nonlinearities. By contrast, mechanisms involving plasticity deeper within the recurrent dynamics become increasingly sensitive to changes in the network operating point. However, if motor-cortical activity is indeed associated with a sufficiently large recurrent-weight variance—which is difficult to determine experimentally—multiple plasticity sites could contribute concurrently to adaptation. Distinguishing their respective contributions would then become essential. Perhaps the clearest experimental test for distinguishing the contributions of each plasticity site would be to evaluate reassociation across multiple levels of neural representation. One should compare the latent factors, the readout activity, and the combined readout and non-readout (i.e., recorded neural activity not used for control, if available) activities. Reaiming predicts systematic reassociation.

## Acknowledgments

We thank Ezekiel Williams for valuable comments on the manuscript. The authors’ research was supported in part by an IVADO postdoctoral fellowship, Canada First Re-search Excellence Fund/Apogée (AP), the Simons Foundation (GL and ALO), a Google Faculty Award (ALO and GL) and an NSERC Discovery Grant (GL). GL further acknowledges support from the Canada CIFAR AI Chair Program and the Canada Re-search Chair in Neural Computations and Interfacing (CIHR, tier 2). ALO further acknowledges support from the Cherng Jia and Elizabeth Yun Hwang endowment to the University of Washington, Seattle. The content of the present paper is solely the responsibility of the authors and does not necessarily represent the official views of the funding agencies.

## A Bias-learning cannot account for adaptation in the model

The bias ***b*** cannot be made the only learnable parameter for adaptation because then

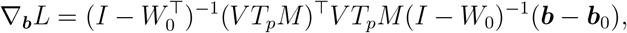

and ***b*** remains fixed at ***b***_0_ during training. The bias—when allowed to vary—can only have an impact on adaptation when either *W* or *Θ* is also plastic. However, bias-learning could have an impact on adaptation when the activation function is nonlinear (Williams et al., 2025).

## B Invariance of ℳ during input-weight learning

We show that the component of *U* in ker {*G*}, denoted ℳ, is invariant under gradient descent. To this end, we decompose the initialization of the input weights *U* as

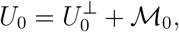

where 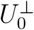 is orthogonal to ker{*G*} whereas ℳ_0_ ∈ ker{*G*}. One gradient update gives

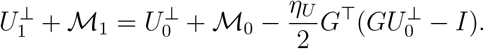

By orthogonality, we identify 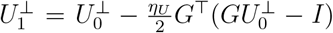 and ℳ_1_ = ℳ_0_. Thus, the component of *U* in ker{*G*} is left unchanged by the update. Iterating this argument shows that gradient descent converges to

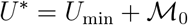

where ℳ_0_ = (*I* − *G*^+^*G*)*U*_0_.

## C Invariance of the gradient with respect to *W* under a rescaling of the input weights

We prove that the loss gradient with respect to *W* is invariant when we rescale the input weight as 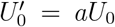 with *a* ≠ 0. Since Σ_0_ depends quadratically on *U*_0_, we have 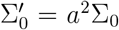 and 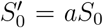. Hence the normalized covariance 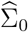 is invariant, implying Λ′ = Λ, *E*′ = *E, T* ′ = *a*^−1^*T* and 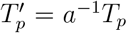. The scaling cancels in the definition of the decoder

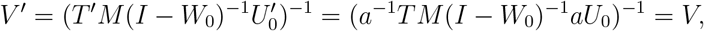

so the network output *Y* and the error ℰ are unchanged. Meanwhile, the neural activity rescales linearly,

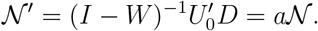

Substituting into the gradient expression shows that ∇_*W*_ *L*′ = ∇_*W*_ *L*. Therefore, when ***b***_0_ = **0**, *σ*_*U*_ can be chosen arbitrarily when analyzing recurrent-weight learning.

## Footnotes

1 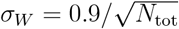 denotes the *N*_tot_ × *N*_tot_ identity matrix. In general, *I*_*n*_ is the identity matrix of size *n*.

2 We note that the bias ***b*** cannot be made the only learnable parameter for adaptation (Appendix A).

3 For a sufficiently small learning rate *η*_*U*_ .

